# Detection of homozygous and hemizygous partial exon deletions by whole-exome sequencing

**DOI:** 10.1101/2020.07.23.217976

**Authors:** Benedetta Bigio, Yoann Seeleuthner, Gaspard Kerner, Melanie Migaud, Jérémie Rosain, Bertrand Boisson, Carla Nasca, Anne Puel, Jacinta Bustamante, Jean-Laurent Casanova, Laurent Abel, Aurelie Cobat

## Abstract

The detection of copy number variations (CNVs) in whole-exome sequencing (WES) data is important, as CNVs may underlie a number of human genetic disorders. The recently developed HMZDelFinder algorithm can detect rare homozygous and hemizygous (HMZ) deletions in WES data more effectively than other widely used tools. Here, we present HMZDelFinder_opt, an approach that outperforms HMZDelFinder for the detection of HMZ deletions, including partial exon deletions in particular, in typical laboratory cohorts that are generated over time under different experimental conditions. We show that using an optimized reference control set of WES data, based on a PCA-derived Euclidean distance for coverage, strongly improves the detection of HMZ deletions both in real patients carrying validated disease-causing deletions and in simulated data. Furthermore, we develop a sliding window approach enabling HMZDelFinder-opt to identify HMZ partial deletions of exons that are otherwise undiscovered by HMZDelFinder. HMZDelFinder_opt is a timely and powerful approach for detecting HMZ deletions, particularly partial exon deletions, in laboratory cohorts, which are typically heterogeneous.

## INTRODUCTION

Copy number variations (CNVs) are unbalanced rearrangements, classically covering more than 50 base pairs (bp), that increase or decrease the number of copies of specific DNA regions (1,2). There is growing evidence to implicate CNVs in common and rare diseases (1,3–5). CNVs have also been linked to adaptive traits, in environmental contexts for example (3). It has been recently estimated that CNVs affect ~5–10% of the genome, suggesting that a number of potentially disease-causing CNVs have yet to be discovered (1,6). Nextgeneration sequencing (NGS) techniques, such as whole-genome and whole-exome sequencing (WGS and WES), provide unprecedent opportunities for studying CNVs. Computational tools using data from WGS have been successfully used to detect CNVs (7–10), but WES-based methods have met with more limited success, mostly due to the nature of targeted enrichment protocols (11–13). Common WGS-based methods use breakpoints, the regions in which the rearrangements occur, to detect CNVs. By contrast, WES focuses on noncontiguous genomic targets (the exons), and most breakpoints are not sequenced. Hence, current WES-based approaches for detecting CNVs use the read depth (or coverage information) as a proxy for copy number information.

The HMZDelFinder algorithm is a recently developed coverage-based method for detecting rare homozygous and hemizygous (HMZ) deletions (14). This subset of CNVs may result in null alleles and a complete loss of gene function. Their identification may, therefore, lead to the discovery of novel genes or variations underlying Mendelian diseases. HMZDelFinder jointly evaluates the normalized per-interval coverage of all the samples of the entire dataset, making it possible to detect rare exonic HMZ deletions while minimizing the number of falsepositive calls due to low-coverage regions. HMZDelFinder outperformed other CNV-calling tools, such as CONIFER (15), CoNVex (16), XHMM (17), ExonDel (18), CANOES (19), CLAMMS (20) and CODEX (21), particularly for the detection of single-exon deletions (i.e. deletions spanning only one exon) (14). However, two major limitations remain to be addressed. First, HMZDelFinder has been optimized to detect HMZ deletions from an entire dataset (>500) of homogeneous exome data. Its performance for typical laboratory cohort, which include exome data generated over time, often under different conditions, is, therefore, not optimal. Second, HMZDelFinder was not designed for the systematic detection of partial exon deletions (i.e. deletions spanning less than one exon). Here, we provide HMZDelFinder_opt, a method that extends the scope of HMZDelFinder by improving the performance of the algorithm for the calling of HMZ deletions in typical laboratory cohorts, which are generated over time, and by allowing the systematic detection of partial exon deletions.

## MATERIALS AND METHODS

### Patient Cohort

The 3,954 individuals used in this study were recruited in collaborations with clinicians, and most of them present different severe infectious diseases. Probands’ family members account for the rest. Although these individuals do not form a random sample, they were ascertained through a number of distinct phenotypes and in different countries. Cohort-specific effects are, therefore, not expected to bias patterns of variation. All study participants provided written informed consent for the use of their DNA in studies aiming to identify genetic risk variants for disease. IRB approval was obtained from The Rockefeller University and Necker Hospital for Sick Children, along with a number of collaborating institutions.

### WES and bioinformatic analysis

WES and bioinformatics analysis were performed as previously described (22). Briefly, genomic DNA was extracted and sheared with a Covaris S2 Ultra-sonicator. An adaptor-ligated library (Illumina) was generated, and exome capture was performed with either SureSelect Human All Exon kits (V5-50Mb, V4-50Mb, V4-71Mb, or V6-60Mb) from Agilent Technologies, or xGen Exome Research 39Mb Panel from Integrated DNA Technologies (IDT xGen). Massively parallel WES was performed on a HiSeq 2000 or 2500 machine (Illumina), generating 100- or 125-base reads. Quality controls were applied at the lane and fastq levels. Specifically, the cutoff used for a successful lane is Pass Filter > 90%, with over 250 M reads for the high-output mode. The fraction of reads in each lane assigned to each sample (no set value) and the fraction of bases with a quality score > Q30 for read 1 and read 2 (above 80% expected for each) were also checked. In addition, the FASTQC tool kit (www.bioinformatics.babraham.ac.uk/projects/fastqc/) was used to review base quality distribution, representation of the four nucleotides of particular k-mer sequences (adaptor contamination). We used the Genome Analysis Software Kit (GATK) (version 3.2.2 or 3.4-46) best-practice pipeline to analyze our WES data(23). Reads were aligned with the human reference genome (hg19), using the maximum exact matches algorithm in Burrows–Wheeler Aligner (BWA)(24). PCR duplicates were removed with Picard tools (picard.sourceforge.net/). The GATK base quality score recalibrator was applied to correct sequencing artifacts.

### Positive controls

The five WES samples used as positive controls carry rare HMZ disease-causing deletions that were confirmed with state-of-the-art molecular approaches (25–27). Specifically, these HMZ deletions comprise one or more exons and have different lengths as follows (SI Table 1). P1 carries a deletion of exons 21 to 23 in *DOCK8* (10,800 bp) that was validated by multiplex ligation-dependent probe amplification (MLPA). The deletion in *DOCK8* was functionally linked to staphylococcus infection (25). P2 had a deletion of exon 5 in *NCF2* (134 bp) that was also validated by MLPA and found to be causal in chronic granulomatous disease (manuscript in preparation). P3’s deletion spanned exons 2 to 8 in *IL12RB1* (13,000 bp) and was validated by sanger sequencing. This deletion was demonstrated to be causal for a Mendelian susceptibility to mycobacterial disease (26). P4 has a deletion of the entire *CYBB* (3,400,000 bp) validated by MLPA and CGH array that resulted in chronic granulomatous disease (27). Finally, P5 is a patient with hyper IgE syndrome carrying a deletion of exons 7 to 15 in entire *DOCK8* (28,000 bp) that was validated by Sanger sequencing. *CYBB* is on the X chromosome while all other genes are autosomal.

### HMZDelFinder-opt

The general workflow used in HMZDelFinder-opt is depicted in SI Figure 1. First, HMZDelFinder_opt computes coverage profiles from the BAM files of the entire dataset. Second, the Principal component analysis (PCA) is calculated from a covariance matrix based on standardized coverage profiles and a k nearest neighbors algorithm is used to select the reference control set. Third, the BAM file of a given sample and the BAM files of the reference control set are used as input of HMZDelFinder to detect HMZ deletions. Fourth, when HMZDelFinder_opt is provided with the parameter -sliding_window_size and the related size, it will employ a sliding window approach for identification of partial deletions of exons. Each of these steps is described in the following paragraphs.

### Principal component analysis (PCA) and k nearest neighbors algorithm

The PCA was performed on the coverage profile of the 3,954 WES using per-exon coverage. Specifically, for each sample, the coverage profile was calculated using the mean depth of coverage of the 194,528 exons from the consensus coding sequences (CCDS) annotation of GRCh37 obtained using biomaRt (28). The PCA was then performed using the ‘prcomp’ function from R 3.5.1 on the scaled coverage profiles. To select the reference control set for a given sample, we computed pairwise weighted Euclidean distances between individuals i and j based on the first 10 principal components from the PCA using the ‘dist’ function of R 3.5.1, using the formula:

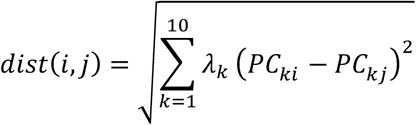

where PC is the matrix of principal components (PCs) calculated on common variants and λ_k_ the eigenvalue corresponding to the k-th principal component PC_k_.

### HMZDelFinder

We used the HMZDelFinder algorithm as described (14). In brief, HMZDelFinder calculates per-exon read depth (reads per thousand base pairs per million reads; RPKM) to detect HMZ deletions. For our purpose of covering all the coding regions, we employed an interval file containing all coding sequences from Gencode. For a given interval, the criteria to call a deletion are as follows: 1) RPKM < 0.65 and 2) frequency of the deletion within the dataset ≤ 0.5%. Filtering criteria at the interval and sample levels include removal of low quality intervals (RPKM median < 7 across all samples) and removal of low quality samples (2% with highest number of calls). When using the optional absence of heterozygosity (AOH) step, HMZDelFinder uses VCF files to filter out deletions not falling in AOH regions, assuming that rare and pathogenic homozygous deletions are likely to be located within larger AOH regions due to the inheritance of a shared haplotype block from both parents. Finally, to prioritize deletions, z-scores are computed. The z-score of a deletion measures the number of standard deviations between the coverage of the deleted interval in a given sample compared to the mean coverage of the same interval in the rest of the dataset. A very low z-score indicates high mean coverage with low variance in the dataset and very low (or no coverage at all) in a given sample. Hence, lower z-scores denote higher confidence in a given deletion.

### Sliding window approach and simulated data

We simulated deletions of variable size in 200 randomly selected individuals among our in-house cohort but excluding the oldest samples (V4-50Mbp capture kit), due to a lower quality than present standards. Two different exons were selected to undergo simulated deletions: a favorable case, exon 11 from LIMCH1 gene (409bp) with a mean coverage of approximately 85X in our samples, and an unfavorable case, exon 4 from RPL15 gene (406 bp) with a mean coverage of 15X in our samples. For both exons, we deleted a segment of 25%, 50%, 75% or 100% of the exon size, using the ‘-v’ argument of the ‘bedtools intersect’ command (bedtools v1.9) on the BAM file to remove all reads overlapping the segment. We then ran HMZDelFinder and HMZDelFinder_opt (with and without the --sliding_windows parameter) on the whole BAM files. Specifically, we applied a sliding window approach, in which each exon was divided into 100 bp windows, with 50 bp overlaps, and BAM files for individual exomes were transformed into per-window read depths. In a separate analysis, we used 50 bp windows, with 25 bp overlaps.

### Analysis of common deletions

To determine whether some of the called deletions were previously reported as common deletions, we utilized the CNVs from the Gold Standard track (hg19 version dated 2016-05-15) of the Database of Genomic Variants (DGV), a highly curated resource that collects CNVs in the human genome (29). We retained only entries with field ‘variant_sub_type’ equal to ‘Loss’ and frequency greater than 1%. We then crossed the retained entries with the deletions called by HMZDelFinder and HMZDelFinder_opt in the positive controls. Deletions were considered common in the DGV database when they overlapped at least 50% with the retained entries from the DGV database.

## RESULTS

### Optimization of the reference control set in HMZDelFinder_opt

We first aimed to improve the performance of HMZDelFinder for detecting HMZ deletions in typical heterogeneous laboratory cohorts, which were generated over time and in different experimental settings (e.g. capture kit). We reasoned that comparing a given sample with an optimized reference control set would limit the impact of the background variability intrinsic to exome data, thereby improving the performance of HMZDelFinder. We designed the optimized reference control set as a selection of samples with similar coverage profiles (SI Figure 1). We did this by first performing a principal component analysis (PCA) of the depth of coverage for consensus coding sequences (CCDS) for 3,954 exomes from our in-house cohort, including mostly patients with severe infectious diseases. As expected, given the different sequencing conditions used for whole-exome sequencing (SI Table 2), the coverage profiles of the samples were highly variable (Figure 1). The first two principal components (PCs) of the PCA identified six distinct clusters, mostly reflecting the capture kit used (Figure 1). Interestingly, two different clusters (clusters 1 and 2 on Figure 1) corresponded to the V4-71Mb capture kit, the difference between these clusters being associated mostly with a minor change in the sequencing chemistry of the kit, leading to a significant improvement in coverage profile for the more recently generated exome data (SI Figure 2). We then used the first 10 PCs to calculate the pairwise weighted Euclidean distances between all samples (30) (see methods). We used this metric to determine, for each sample of interest, the closest neighbors, for use as the reference control set in HMZDelFinder_opt.

**Figure 1:**
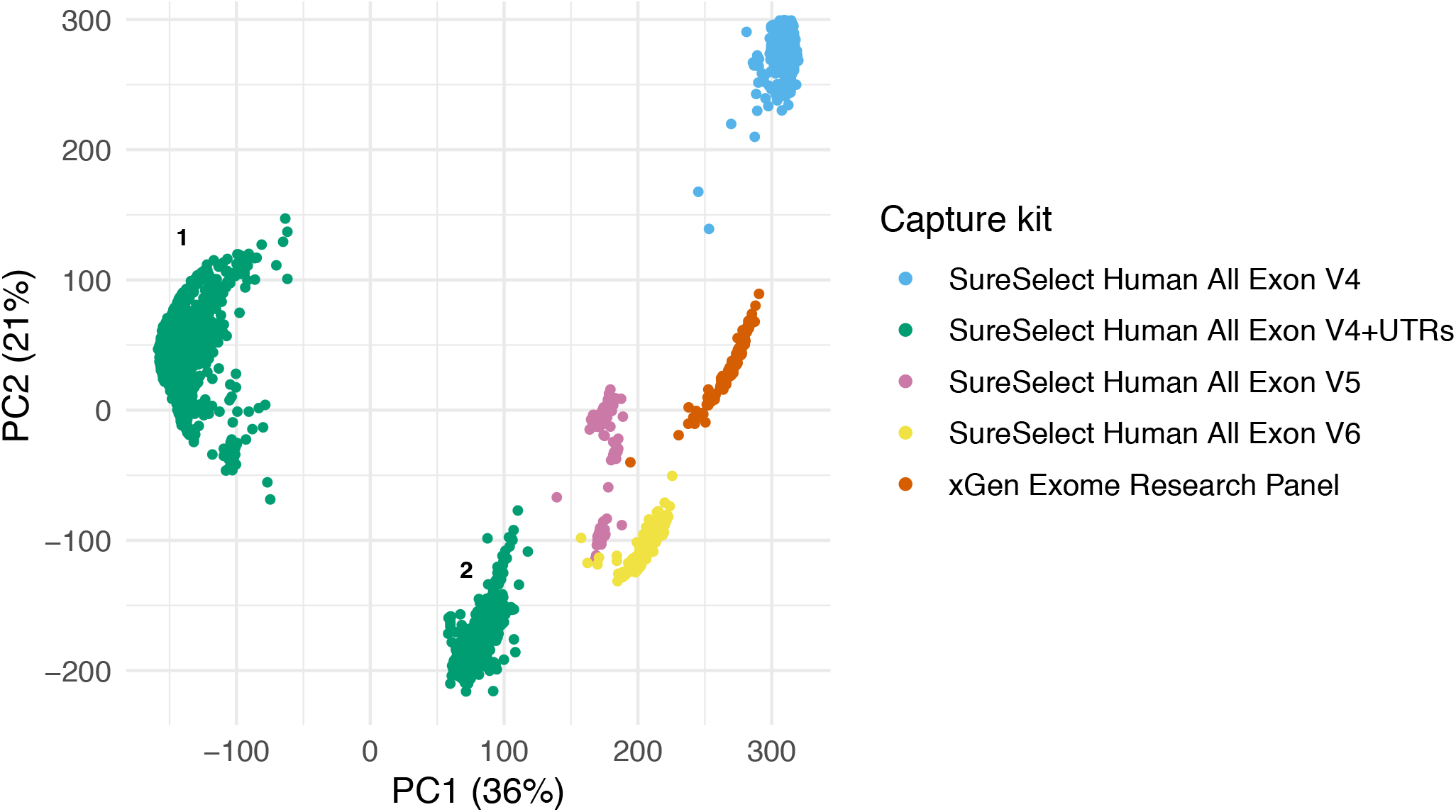
Principal Component Analysis (PCA) of the WES coverage. The PCA was computed from the coverage profiles of consensus coding sequences (CCDS) from 3,954 individuals. Dots are color-coded by the type of the capture kit used for sequencing.). Two different clusters (clusters 1 and 2) corresponded to the V4-71Mb capture kit. See also SI Figure 2.

We then compared the performances of HMZDelFinder_opt and HMZDelFinder, using five WES samples carrying validated rare HMZ disease-causing deletions of different lengths as positive controls (SI Table 1, methods). Specifically, we tested the ability of HMZDelFinder_opt and HMZDelFinder to detect the validated deletions, and we also compared the total numbers of deletions called and their *z*-scores (see Methods). In HMZDelFinder_opt, we compared reference control sets of different size (ranging from 50 to 500, SI Figure 3), selected for each sample as described above. In HMZDelFinder, we used the entire dataset, consisting of 3,954 WES samples. For both approaches, the final set of called deletions for each sample was narrowed down to the capture kit corresponding to the patient WES data. We chose to benchmark HMZDelFinder because it has been shown to perform at least as well as, and sometimes better than several widely used and actively maintained detection tools (14).

Both HMZDelFinder and HMZDelFinder_opt successfully detected all five confirmed HMZ deletions in the positive controls, regardless of the size of the reference control set (Table 1). However, HMZDelFinder_opt detected a smaller total number of deletions than HMZDelFinder (Table 1). Specifically, the total number of deletions ranged from one to 21 deletions for HMZDelFinder_opt, and from 11 to 2,586 for HMZDelFinder, suggesting that a smaller number of false-positive calls were obtained with HMZDelFinder_opt. Using the optional filtering step based on the absence of heterozygosity (AOH) information for HMZDelFinder (see methods) decreased the number of deletions detected, but this number nevertheless remained much higher than that for HMZDelFinder_opt (Table 1). We hypothesized that the large difference between the two methods for P1 reflected the low quality of exome data for this patient. Indeed, the mean coverage and the proportion of bases with coverage above 10x were much lower for P1 than for the other four patients (e.g. only 68.9% of bases had a coverage above 10x for P1, versus >99% for the other patients) (SI Table 1), leading to a large number of likely false positive deletions detected when not using an appropriate reference control set with similar coverage. Consistently, the number of deletions detected for P1 with HMZDelFinder_opt was larger with the largest reference sample size (500) (Table 1). We therefore performed subsequent HMZDelFinder_opt analyses with a reference sample size of 100, which provided a good compromise between the algorithm performance and computation time.

**Table 1:** Comparison of the results between HMZDelFinder_opt and HMZDelFinder by using five positive controls carrying validated rare HMZ disease-causing deletions. Both HMZDelFinder_opt and HMZDelFinder (with or without AOH filtering step) detect the confirmed deletions. HMZDelFinder_opt detects a lower number of other deletions and ranks higher the confirmed deletion as compared to HMZDelFinder with or without AOH filtering step.

|  |  | P1 | P2 | P3 | P4 | P5 |
| --- | --- | --- | --- | --- | --- | --- |
| KIT |  | V4-50MB | V6-60MB | V5-50MB | V5-50MB | V6-60MB |
| METHOD | N NEIGHBORS | Confirmed deletion (Rank/Total number of deletions) |  |  |  |  |
| HMZDeIFinder_opt | 50 | DOCK8<br>(1/11) | NCF2 (1/2) | IL12RB1<br>(1/1) | CYBB<br>(3/5) | DOCK8<br>(1/3) |
|  | 100 | DOCK8<br>(1/11) | NCF2 (1/2) | IL12RB1<br>(1/1) | CYBB<br>(4/5) | DOCK8<br>(1/2) |
|  | 200 | DOCK8<br>(1/11) | NCF2 (1/3) | IL12RB1<br>(1/1) | CYBB<br>(4/5) | DOCK8<br>(1/3) |
|  | 500 | DOCK8<br>(4/21) | NCF2 (1/2) | IL12RB1<br>(1/3) | CYBB<br>(3/5) | DOCK8<br>(1/2) |
| HMZDeIFinder | All | DOCK8<br>(1/2586) | NCF2<br>(120/120) | IL12RB1<br>(4/11) | CYBB<br>(7/13) | DOCK8<br>(1/163) |
| HMZDeIFinder AOH | All | DOCK8<br>(1/457) | NCF2 (37/37) | IL12RB1<br>(2/5) | CYBB<br>(4/7) | DOCK8<br>(1/46) |

We then compared the rankings of the confirmed deletions between the two algorithms, using the *z*-score provided by HMZDelFinder (see method). While the two approaches ranked the confirmed disease-causing deletions for P1 and P5 first, HMZDelFinder_opt ranked higher the confirmed disease-causing deletions for P2, P3 and P4 than HMZDelFinder (Table 1; Figure 2). Moreover, *z*-scores were consistently better with HMZDelFinder_opt (Figure 2) than with HMZDelFinder, leading to a more specific discovery of true HMZ deletions. Again, using the AOH option for HMZDelFinder slightly improved the ranking (Table 1), but did not change the *z*-score ranking. Together, these results suggest that HMZDelFinder_opt gives better z-scores for deletions than HMZDelFinder, which should lead to higher sensitivity in the general case.

**Figure 2:**
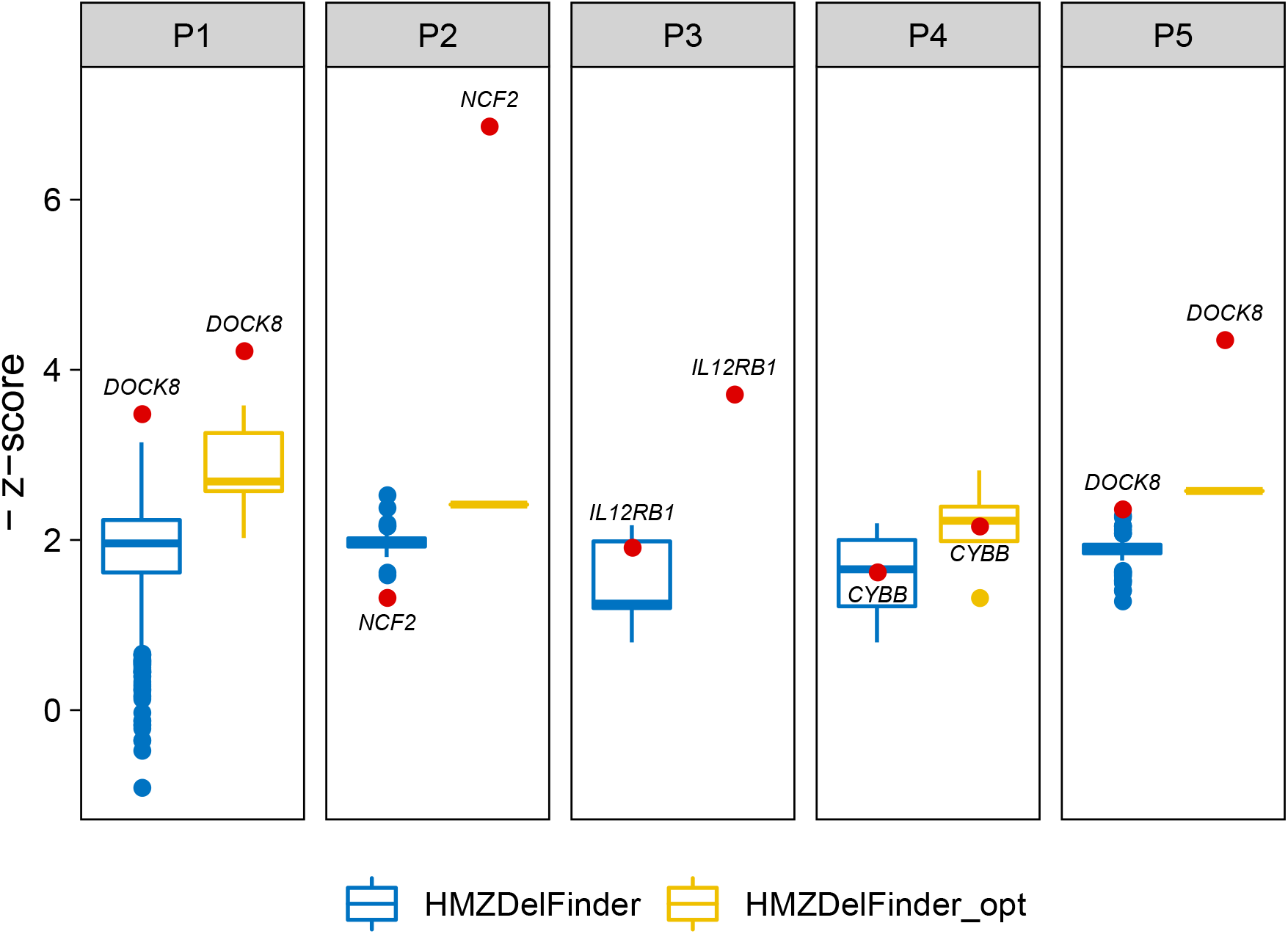
Comparison of the ranking of the deletions called by HMZDelFinder_opt and HMZDelFinder in five positive controls carrying validated rare HMZ disease-causing deletions. The ranking is expressed as - z-score. Lower z-scores (and higher ranking) indicate more confidence in a given deletion. The confirmed deletions ranked 1^st^ in P1, P2, P3, P5 with HMZDelFinder_opt while they ranked 1^st^ only in P1 and P5 with HMZDelFinder as shown by the red dots in the blue (HMZDelFinder) and yellow (HMZDelFinder_opt) distributions. The ranking was consistently higher with HMZDelFinder_opt than with HMZDelFinder. Results are shown for HMZDelFinder_opt using 100 as size of the reference control set.

Finally, we studied the HMZ deletions called by both approaches, in addition to the validated ones, to determine whether some of the deletions identified were reported as common deletions. We used the CNVs from the gold standard track of the Database of Genomic Variants (DGV), a highly curated resource containing CNVs from the human genome (29). We focused on the positive controls with high data quality (P2, P3, P4 and P5), and found that the HMZ deletions called by HMZDelFinder_opt were more enriched in common deletions (frequency > 1%) than those called by HMZDelFinder (SI Table 3). Among the 6 and 303 additional HMZ deletions called by HMZDelFinder–opt (with the reference control set of 100 exomes) and HMZDelFinder, 50% and 1%, respectively, were present in the DGV database (SI Table 3), suggesting that the deletions called by HMZDelFinder_opt were enriched in true deletions. Overall, these findings demonstrate that the use of an appropriate reference control set of WES data based on a PCA-derived coverage distance improves the performance of HMZDelFinder. These results also provided a first validation of HMZDelFinder_opt for five confirmed disease-causing HMZ deletions.

### Detection of HMZ partial exon deletions by HMZDelFinder_opt

In HMZDelFinder, individual exome BAM files are transformed into per-exon read depths, facilitating a more efficient detection of single-exon HMZ deletions than can be achieved with other classical CNV-calling algorithms (14). Here, we aimed to address the need for the identification of even smaller HMZ deletions, spanning less than an exon (partial exon deletions). To this end, we used HMZDelFinder_opt with a sliding window approach, in which each exon was divided into 100 bp windows, with 50 bp overlaps, and BAM files for individual exomes were transformed into per-window read depths. We tested this approach by simulating deletions in two exons of similar size (~400 bp) but with different mean coverages in a randomly selected dataset of 200 WES samples from our in-house cohort. The deletions spanned 100%, 75%, 50% or 25% of either exon 11 of *LIMCH1* (409 bp, ~85x mean coverage) or exon 4 of *RPL15* (406 bp, ~15x mean coverage). We used these datasets to compare the performances of HMZDelFinder_opt with sliding windows of 100 bp (HMZDelFinder_opt+sw100), HMZDelFinder_opt without sliding windows (HMZDelFinder_opt), and the original HMZDelFinder. For HMZDelFinder_opt+sw100 and HMZDelFinder_opt, we used reference control sets of size 100.

For deletions spanning the full exon (100%), we confirmed that HMZDelFinder_opt had a detection rate (98% and 93% for exons with higher and lower coverage, respectively; Figure 3) similar to that of HMZDelFinder (98% and 93% for exons with higher and lower coverage, respectively). However, the total number of HMZ deletions called by HMZDelFinder_opt was only one eighth the total number of HMZ deletions called by HMZDelFinder (median number of HMZ deletions: 2 vs. 13 SI Figure 4). The detection rate was slightly higher when sliding windows were used (detection rate for HMZDelFinder_opt+sw100 of 99% and 94% for exons with a higher and lower coverage, respectively), but at the cost of a slightly larger total number of HMZ deletions called than for HMZDelFinder_opt (median number of deletions: 5 vs. 2). Nevertheless, the total number of HMZ deletions called by HMZDelFinder_opt+sw100 remained lower than the total number of HMZ deletions called by HMZDelFinder.

**Figure 3:**
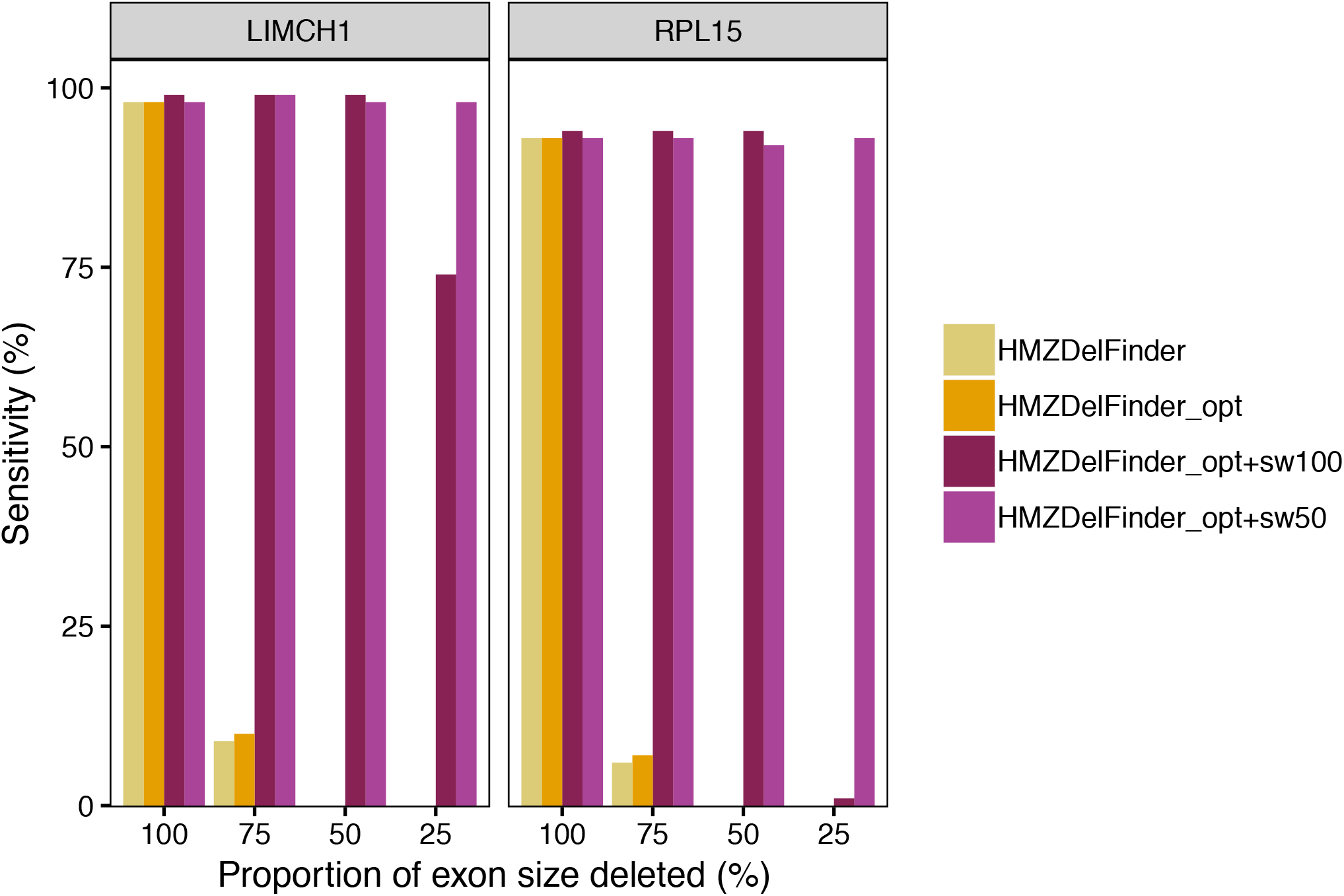
Comparison of HMZDelFinder_opt with or without sliding windows and HMZDelFinder by using simulated data. Proportions of deletions detected in simulated data in the higher (LIMCH1) or lower (RPL15) covered exons by using HMZDelfinder (yellow), HMZDelFinder_opt (orange), HMZDelFinder_opt+sw100 (red), HMZDelFinder_opt+sw50 (pink).

For partial exon deletions, the detection rates of HMZDelFinder and HMZDelFinder_opt were much lower, at less than 10% for deletions spanning 75% of the exon and 0% for deletions spanning 25% or 50% of the exon. Conversely, HMZDelFinder_opt+sw100 succeeded in detecting simulated deletions spanning 50% or 75% (200 bp or ~300 bp) of both exon 11 of *LIMCH1* and exon 4 of *RPL15* in 99% of the samples, with a median number of called HMZ deletions of 5 (Figure 3, SI Figure 4). For deletions spanning 25% of the exon (~100 bp), HMZDelFinder_opt+sw100 had a detection rate of 74% for the exon with the highest coverage in *LIMCH1*, but it failed to detect the deletions in the exon with the lowest coverage in *RPL15*. We assessed the performance of this method further, using a smaller sliding window of 50 bp in size, and a step size of 25 bp, to improve granularity. We found that the use of smaller sliding windows with HMZDelFinder_opt+sw50 greatly increased the detection rate for deletions spanning 25% of the exon with the lowest coverage, exon 4 of *RPL15* (93% for sw50 vs. 1% for sw100) and of the exon with the highest coverage in *LIMCH1* (98% for sw50 vs. 74% for sw100) (Figure 3). Thus, the use of a sliding window makes it possible to detect HMZ partial exon deletions that would otherwise be missed, and the use of simulated data further validated HMZDelFinder_opt.

### DISCUSSION

WES offers unprecedent opportunities for identifying HMZ deletions as novel causal determinants of human diseases, but it poses a number of computational challenges. Most current methods for detecting HMZ deletions compare the depth of coverage between a given exome and the rest of the exomes in the dataset. However, coverage depth is heavily dependent on sequencing conditions, which are continually evolving in typical laboratory settings. Thus, the exome data generated over time are inevitably heterogeneous, complicating the discovery of deletions. Using HMZDelFinder_opt with both validated disease-causing deletions and simulated data, we demonstrated that the *a priori* selection of a reference control set with a coverage profile similar to that of the WES sample studied reduced the number of deletions detected, while improving the ranking of the true HMZ deletion. These results are consistent with a recent report showing that the selection of an appropriate reference control set with multidimensional scaling significantly improves the sensitivity of various CNV callers (31). In further support for our findings, the ranking of the known deletion and the number of additional deletions detected by HMZDelFinder_opt start worsening with increasing numbers of controls in the reference set, including neighbors with a less similar coverage profile, as illustrated, for P1, in SI Fig. 3A.

In addition to providing an optimized tool for detecting deletions in typical laboratory cohorts, HMZDelFinder_opt also fills the gap in the study of deletions spanning less than an exon, by providing the first tool for the systematic identification of partial exon deletions. Existing CNV callers are optimized for the detection of either large deletions (usually spanning more than three exons), or deletions of full single exons (14,32). Other established callers, such as GATK, are not designed to detect CNVs and can therefore identify deletions of only a few dozen base pairs (typically up to 50 bp, https://gatkforums.broadinstitute.org/gatk/discussion/5938/using-gatk-tool-how-long-insertion-deletion-could-be-detected and (33)). The human genome contains ~235,000 exons, about 20% of which are larger than 200 bp (34). HMZDelFinder_opt therefore makes possible the systematic discovery of currently unknown HMZ deletions in ~47,000 exons that are not detectable with other tools. In sum, we describe HMZDelFinder_opt, a method for improving the detection of HMZ deletions in heterogeneous exome data that can be used to identify partial exon deletions that would otherwise be missed, through an extension of the scope of HMZDelFinder.

## DATA AVAILABILITY

The code for the PCA-based selection and sliding window is available in the GitHub repository (https://github.com/casanova-lab/HMZDelFinder_opt/).

## ACKNOWLEDGEMENT

We thank the members of the Human Genetics of Infectious Diseases Laboratory for helpful discussions. We also thank Yelena Nemiroskaya, Dominick Papandrea, Mark Woollett, Dana Liu (St. Giles Laboratory of Human Genetics of Infectious Diseases, Rockefeller Branch, The Rockefeller University, New York, New York, USA), and Cécile Patissier, Lazaro Lorenzo-Diaz, Christine Rivalain (Laboratory of Human Genetics of Infectious Diseases, Necker Branch, INSERM U1163, Necker Hospital for Sick Children, Paris, France) for their assistance.

