## Supplementary information for "Detection of homozygous and hemizygous partial exon deletions by whole-exome sequencing"

Benedetta Bigio[1,2,3], Yoann Seeleuthner[2,3], Gaspard Kerner[2,3], Melanie Migaud[2,3], Jérémie Rosain[2,3], Bertrand Boisson[1,2,3], Carla Nasca[4], Anne Puel[2,3], Jacinta Bustamante[1,2,3,5], Jean-Laurent Casanova[1,2,3,6,7], Laurent Abel[1,2,3,#,*], Aurelie Cobat[2,3,#,*]

[1]St. Giles Laboratory of Human Genetics of Infectious Diseases, Rockefeller Branch, The Rockefeller University, New York, NY 10065, USA

[2]Laboratory of Human Genetics of Infectious Diseases, Necker Branch, INSERM U1163, Necker Hospital for Sick Children, 75015 Paris, France

[3]University of Paris, Imagine Institute, 75015 Paris, France

[4]Laboratory of Neuroendocrinology, The Rockefeller University, New York, NY 10065, USA

[5]Study Center of Immunodeficiencies, Necker Hospital for Sick Children, 75015 Paris, France

[6]Pediatric Hematology-Immunology Unit, Necker Hospital for Sick Children, 75015 Paris, France

[7]Howard Hughes Medical Institute, New York, NY 10065, USA

[#] indicates equal contributions

* To whom correspondence should be addressed.

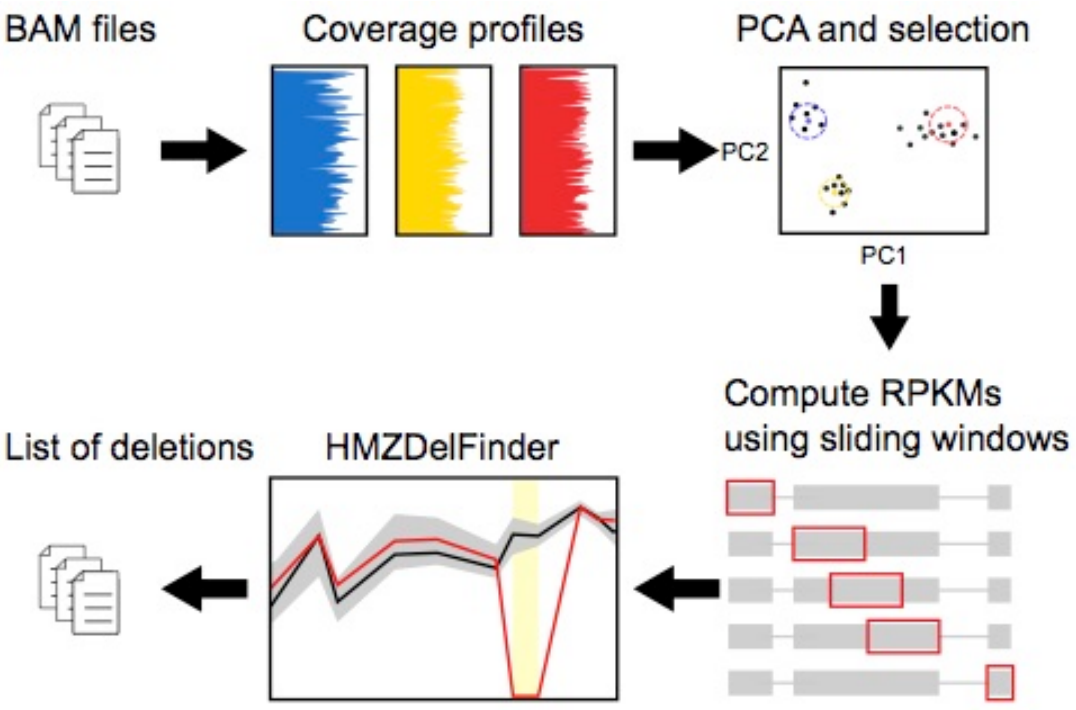


**SI Figure 1: Schematic representation of the method employed by HMZDelFinder_opt to detect partial-exon homozygous and hemizygous deletions.** First, HMZDelFinder_opt computes coverage profiles from the BAM files. The PCA is then calculated from a covariance matrix based on standardized coverage profiles and a k nearest neighbors algorithm is used to select the reference control set. The BAM file of a given sample and the BAM files of the reference control set are used as input of HMZDelFinder to detect homozygous and hemizygous deletions. In addition, HMZDelFinder_opt accepts a parameter (-sliding_window_size) to employ a sliding window approach for identification of partial-exon deletions.

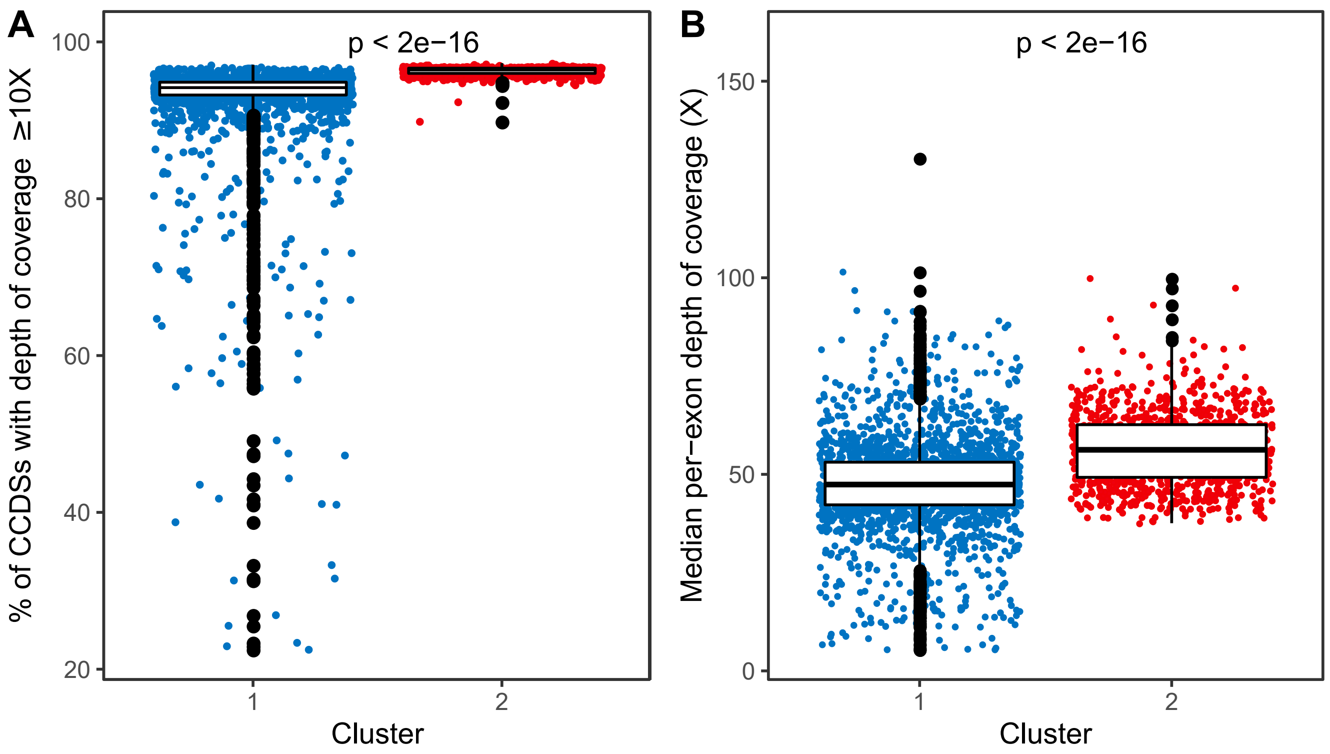


**SI Figure 2: Coverage in the two exome clusters revealed by PCA within the exomes generated by the V4-71Mb capture kit.** Both the number of CCDSs with at least 10X (A) and the depth of coverage per exon (B) are significantly higher ($p<2^{-16}$) in the most recent V4-71Mb exomes (Cluster 2) than in the oldest V4-71Mb exomes (Cluster 1). Effect size: A, Cohen's d=0.45; B, Cohen's d=0.8.

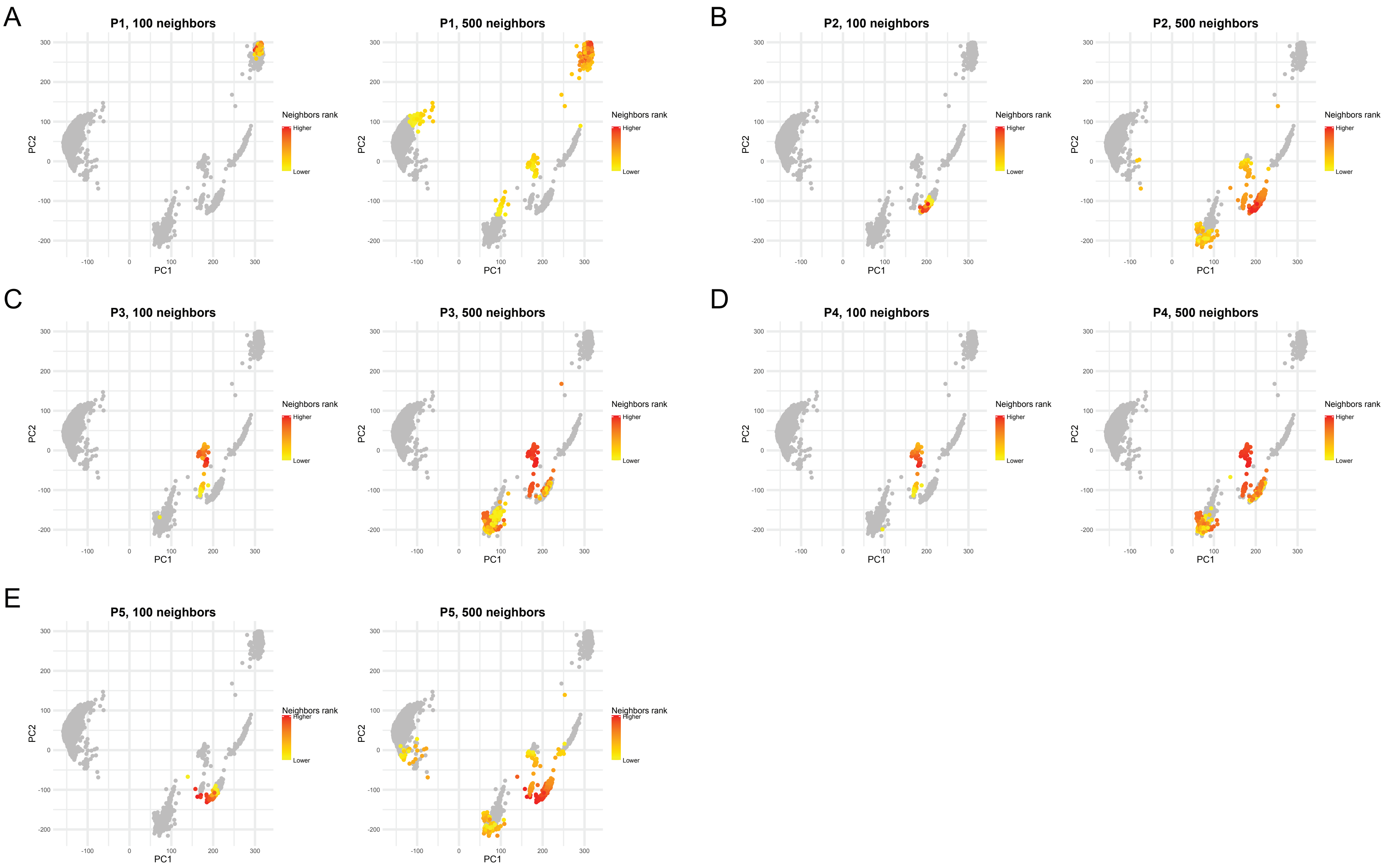


**SI Figure 3: Closest neighbors of the positive controls as function of the size of the reference control set.** A total of 100 and 500 neighbors are showed for P1 (A), P2 (B), P3 (C), P4 (D), and P5 (E).

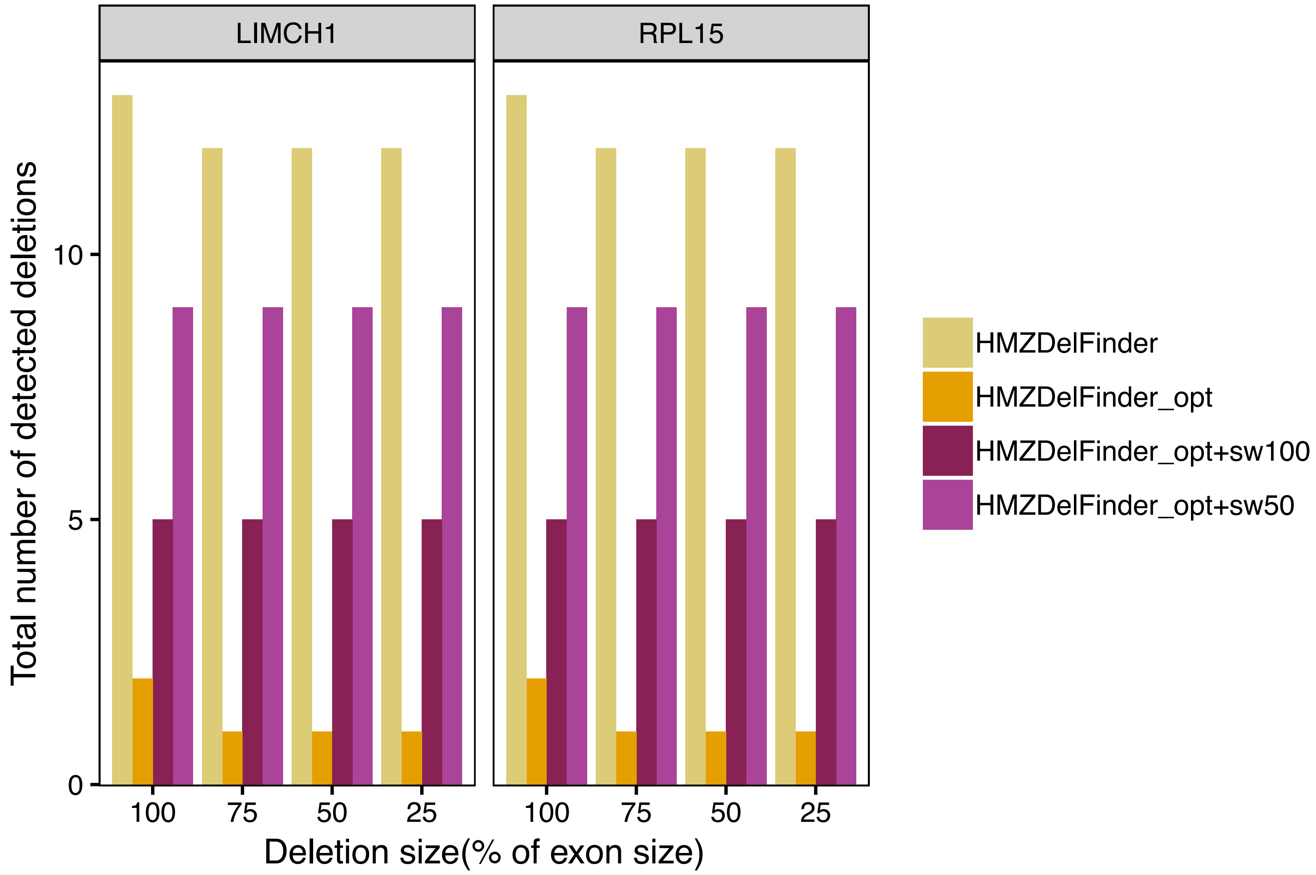


**SI Figure 4**: Median number of detected deletions in the simulated data in the higher (LIMCH1) or lower (RPL15) covered exons by using HMZDelfinder (yellow), HMZDelFinder_opt (orange), HMZDelFinder_opt+sw100 (red), HMZDelFinder_opt+sw50 (pink).

| Patient | Confirmed Homozygous Deletion | | | | Exome | |
|---|---|---|---|---|---|---|
| | Location | Gene | Size (kbp) | Validation method | Mean Coverage | % Bases above 10% |
| **P1** | Chr 9, Exons 21 to 23 | DOCK8 | 10.8 | MLPA | 23 | 68.9 |
| **P2** | Chr 1, Exon 5 | NCF2 | 0.13 | MLPA | 115 | 99.5 |
| **P3** | Chr 19, Exons 2 to 8 | IL12RB1 | 13 | Sanger sequencing | 206 | 99.5 |
| **P4** | Chr X, Whole gene | CYBB | 3,400 | MLPA and CGH array | 156 | 99.2 |
| **P5** | Chr 9, Exons 7 to 15 | DOCK8 | 28 | Sanger sequencing | 66 | 99.5 |

**SI Table 1:** Validated rare HMZ disease-causing deletions and exome coverage in the five exomes used as positive controls.

| Kit | Kit (full name) | Number (Percentage) of Exomes | Median Coverage (SD) | % bases above 10X |
|---|---|---|---|---|
| IDT-xGen | xGen Exome Research Panel v2 from Integrated DNA Technologies | 188 (4.8%) | 41.7 (9.5) | 91.4 |
| V4-50Mb | Agilent SureSelect Human All Exon V4 | 354 (9.0%) | 50.0 (15.5) | 83.2 |
| V4-71Mb | Agilent SureSelect Human All Exon V4+UTRs | 3095 (78.3%) | 47.4 (10.2) | 81.0 |
| V5-50Mb | Agilent SureSelect Human All Exon V5 | 101 (2.6%) | 72.4 (43.7) | 70.3 |
| V6-60Mb | Agilent SureSelect Human All Exon V6 | 216 (5.5%) | 125.9 (38.6) | 99.0 |

**SI Table 2:** Distribution of the capture kit in the 3,954 exomes and corresponding coverage metrics.

| | | P2 | P3 | P4 | P5 | TOTAL |
|---|---|---|---|---|---|---|
| | KIT | V6-60MB | V5-50MB | V5-50MB | V4-71MB | |
| METHOD | N NEIGHBORS | (COMMON DELETIONS/NUMBER OF OTHER DETECTED DELETIONS) | | | | |
| HMZDelFinder_opt | 50 | 0/1 (0%) | 0/0 (-) | 2/4 (50%) | 2/2 (100%) | 4/7 (60%) |
| | 100 | 0/1 (0%) | 0/0 (-) | 2/4 (50%) | 1/1 (100%) | 3/6 (50%) |
| | 200 | 0/2 (0%) | 0/0 (-) | 2/4 (50%) | 2/3 (67%) | 4/9 (44%) |
| | 500 | 0/1 (0%) | 0/2 (0%) | 2/4 (50%) | 1/1 (100%) | 3/8 (38%) |
| HMZDelFinder | all | 0/119 (0%) | 0/10 (0%) | 1/12 (8%) | 2/162 (1%) | 3/303 (1%) |

**SI Table 3:** Number and percentage of common deletions (>1% frequency) among the detected deletions (other than the confirmed deletion)
